# RBOHC-Generated ROS Tune GNOM-Dependent Root Halotropism in *Arabidopsis*

**DOI:** 10.64898/2026.01.25.701553

**Authors:** Amir Cohen, Matan Franko, Yvonne Kiere, Yonatan Wexler, Doron Shkolnik

## Abstract

Halotropism—the directional growth of roots away from saline environments—requires coordinated integration of tropic cues. We show that halotropic bending in *Arabidopsis thaliana* roots is fine-tuned by a spatially confined, symmetric reactive oxygen species (ROS) domain generated by the NADPH oxidase RBOHC in elongation-zone epidermal cells. This domain, visualized by dihydrorhodamine-123 staining and confocal microscopy, emerges during the first hours of halostimulation and limits excessive curvature. Reducing ROS, either chemically with ascorbate or diphenyleneiodonium, or genetically in *rbohC* mutants, enhances halotropic bending, whereas *miz2*, defective in the ARF-GEF GNOM, exhibits negative halotropism due to an expanded and mislocalized ROS domain that disrupts spatial restriction. The *miz2 rbohC* double mutant shows a much weaker halotropic response than *rbohC* alone and similarly lacks the halotropic ROS signals in the elongation zone, indicating that GNOM acts upstream of RBOHC-mediated ROS production. Comparisons with hydrotropism—a moisture-seeking response also involving defined ROS distribution—suggest that GNOM-dependent regulation of RBOHC constitutes a shared module for adjusting root orientation to environmental gradients. Understanding these molecular mechanisms is essential for enhancing crop resilience to soil salinity, particularly in the context of increasing soil salinization driven by climate change.

## Introduction

Soil salinization, exacerbated by unsustainable irrigation practices, natural processes, and climate change, leads to the accumulation of salts in agricultural soils and generates heterogeneous salt distribution (Hassani, Azapagic & Shokri 2021). Rising soil salinity severely limits crop productivity and poses a growing threat to global food security (FAO, 2025). Understanding how plants perceive and respond to salt stress—through ion transport, hormone signaling, and reactive oxygen species (ROS) dynamics—is critical for developing strategies to enhance salt tolerance (Munns & Tester 2008). Insights into these molecular mechanisms can inform the design of crops capable of thriving in saline and variable environments, thereby supporting sustainable agriculture now and in the future.

Plants employ several complementary strategies to cope with salinity (Munns 2002; Zhu 2002; Munns & Tester 2008). First, ionic homeostasis mechanisms help prevent toxic cytosolic accumulation of sodium ions (Na⁺), a primary ionic stress factor under saline conditions. These include Na⁺ extrusion via the plasma-membrane antiporter SOS1 (Shi, Ishitani, Kim & Zhu 2000), Na⁺ retrieval by HKT1 transporters (Rus *et al*. 2001; Mäser *et al*. 2002; Sunarpi *et al*. 2005; Shkolnik-Inbar, Adler & Bar-Zvi 2013; Chandran *et al*. 2024), and vacuolar sequestration mediated by NHX antiporters, all contributing to the maintenance of a favorable K⁺/Na⁺ ratio (Apse et al., 1999; Khan, Ali & Yun 2018). In parallel, plants engage in osmotic adjustment through the accumulation of compatible solutes such as proline, glycine betaine, and soluble sugars, which stabilize proteins and membranes and support turgor under osmotic stress (Flowers & Colmer, 2008). Salinity also perturbs cellular redox balance; accordingly, plants reinforce ROS-regulation pathways, modulating both ROS production and scavenging capacity to prevent oxidative damage and maintain metabolic function (Miller, Shulaev & Mittler 2008; Baxter, Mittler & Suzuki 2014; Mittler 2017). Beyond these cellular strategies, plants activate organ-level adaptive responses that reshape root-system architecture. These include reduced primary root elongation, altered lateral root formation, and directional growth responses that help roots navigate heterogeneous saline environments (Galvan-Ampudia & Testerink 2011). One such response, root halotropism, enables roots to actively grow away from localized salt sources and represents a key adaptive mechanism under patchy soil salinity (Galvan-Ampudia et al. 2013; van den Berg et al., 2016).

Halotropism is controlled by a combination of auxin-dependent and auxin-independent mechanisms. Seminal studies have demonstrated that halotropic curvature relies on the establishment of asymmetric auxin distribution across the root tip mediated by PIN2- and AUX1-driven transport (Galvan-Ampudia *et al*. 2013; van den Berg *et al*. 2016; Korver *et al*. 2020). More recently, Zheng et al. (2024) showed that the transcription factor SOMBRERO (SMB) modulates AUX1 expression to generate the auxin imbalance necessary for late-stage halotropic bending, reinforcing the central role of auxin transport in shaping directional growth. Complementing this auxin-based framework, Yu et al. (2022) identified an early, auxin-independent pathway in which salt stress triggers abscisic acid (ABA)-dependent phosphorylation of the microtubule-associated protein SP2L by SnRK2.6, promoting microtubule reorientation and anisotropic expansion in the transition zone. Together, these findings position halotropism as a multilayered process integrating auxin transport and ABA–microtubule signaling to drive root reorientation away from saline zones.

Insights from hydrotropism—the reorientation of roots toward regions of higher water potential—provide a strategic framework for identifying candidate regulators of halotropism. Hydrotropic growth depends on root-tip perception and elongation-zone modulation and often overcomes competing gravitropic cues (Wexler, Schroeder & Shkolnik 2024). Genetic analyses of hydrotropism have revealed only two loci whose loss completely abolishes the response: MIZU-KUSSEI1 (MIZ1) and GNOM/MIZU-KUSSEI2 (MIZ2) (Kobayashi *et al*. 2007; Miyazawa *et al*. 2009). While *miz1* mutants display normal halotropic bending (Yu *et al*. 2022), the potential involvement of GNOM/MIZ2 in halotropism has not yet been explored. Notably, hydrotropic bending occurs independently of auxin and asymmetric auxin distribution (Kaneyasu *et al*. 2007; Shkolnik, Krieger, Nuriel & Fromm 2016), emphasizing mechanistic distinctions between halotropism and hydrotropism. Nevertheless, the two tropic responses may share key regulatory components, such as GNOM, reflecting a common strategy by which roots integrate environmental signals to modulate elongation and counteract gravitropic growth. Together, these observations justify investigating hydrotropism-associated genes as potential mediators of halotropism.

Respiratory burst oxidase homologs (RBOHs) are NADPH oxidases that generate apoplastic ROS, which function as versatile secondary messengers in root development and environmental signaling (Suzuki et al. 2011; Mittler et al., 2022). Among the *Arabidopsis thaliana* isoforms, RBOH protein C (RBOHC; also known as RHD2) is strongly expressed in root epidermal cells, including those in the elongation and differentiation zones of the primary root, where it contributes to spatially localized ROS production (Foreman et al. 2003; Monshausen et al., 2007). Loss-of-function mutants in RBOHC display defective root-hair growth due to impaired ROS generation, highlighting the importance of RBOHC-mediated ROS in expanding root cells (Foreman *et al*., 2003). ROS generated by RBOHC have also been implicated in modulating differential growth during gravitropism and hydrotropism, acting as integrators that balance multiple directional cues to fine-tune tropic outputs without directly driving curvature (Krieger et al., 2016). Despite their established roles in root-hair formation, lateral root emergence, and other tropic processes, the involvement of ROS in halotropism has not yet been investigated.

*Arabidopsis* serves as a powerful model for dissecting the molecular mechanisms that govern root tropic responses and environmental adaptation (Muthert et al., 2020; Wexler, Schroeder & Shkolnik 2024). Studies in *Arabidopsis* have revealed how root bending in gravitropism, hydrotropism, and halotropism is orchestrated through coordinated signaling networks, including auxin transport, ROS dynamics, and Ca²⁺-dependent pathways. Key components of these pathways include the auxin efflux carrier PIN2, the hydrotropism regulator MIZ1, and NADPH oxidases such as RBOHC, which produce the spatially localized ROS required for differential growth during tropic responses (Kobayashi *et al*. 2007; Galvan-Ampudia *et al*. 2013; Krieger *et al*. 2016; Shkolnik, Nuriel, Bonza, Costa & Fromm 2018; Yu *et al*. 2022; Zhang *et al*. 2025). Insights from these studies demonstrate that the interplay between auxin, ROS, and other signaling modules fine-tunes root growth, enabling plants to navigate complex environments. Understanding these mechanisms in the context of halotropism provides a framework for uncovering molecular strategies to enhance plant resilience to soil salinity and to support crop productivity under increasingly saline conditions.

## Materials and Methods

### Plant Material and Growth Conditions

*Arabidopsis thaliana* (ecotype Col-0) and its mutant lines *miz2* (*gnom*^G2852A^) (Miyazawa *et al*. 2009), *rbohC (rhd2)* and *rbohD* (Miller *et al*. 2009) were used in this study. Seeds were surface-sterilized and cold-stratified as described previously (Shkolnik & Bar-Zvi 2008). Germination and seedling growth were carried out on 0.25× Murashige and Skoog (MS) medium (Murashige & Skoog 1962) solidified with 0.7% (w/v) agar. Square Petri dishes were positioned vertically to allow roots to grow along the medium surface. To obtain seedlings with uniformly straight primary roots and ensure reproducible halotropic responses, seedlings were grown partially embedded in the solidified medium, as previously described (Wexler *et al*. 2025). Growth conditions were 16-h light/8-h dark cycles at 22°C. *miz2 rbohC* double mutants were generated by crossing the single mutant lines and selfing the F1 progeny; segregating F2 plants were screened to identify individuals carrying both mutant alleles, and homozygous double mutants were confirmed in the F3 generation. Genotyping of *miz2* and *rbohC* alleles was performed by polymerase chain reaction (PCR) using the following primer pairs (5′→3′): MIZ2-Fwd, CTGTTGATGAACCCGTTCTTGC; MIZ2-Rev, GCAAGCTGCAACAAAGATTCAGC; RBOHC-Fwd, CAACTTTGGTGTTAGGATGATCCATAGCTATACG; RBOHC-Rev, CTTCCCGAAAGTCTTGATTGATGGTCC. PCRs were carried out using Q5 High-Fidelity DNA Polymerase (Takara, Japan) according to the manufacturer’s instructions, with an annealing temperature of 60°C. For *miz2*, the PCR product was subjected to Sanger sequencing to identify the G2852A point mutation. For *rbohC*, presence of the insertion was inferred from the absence of a PCR product, whereas clear amplification was observed in the wild-type.

### Halostimulation System under Continuous Clinorotation

To expose roots to a lateral NaCl gradient (halostimulation) while minimizing gravitropic influence, we adapted the split-agar method originally developed for hydrostimulation by replacing sorbitol with NaCl (Takahashi, Goto, Okada & Takahashi 2002; Krieger *et al*. 2016; Yu *et al*. 2022). Four- or five-day-old seedlings were positioned on control agar-solidified medium occupying half of a 90 mm Petri dish, with root tips placed 5 mm from the medium edge. The other half of the plate was filled with medium supplemented with NaCl (200 mM), allowing gradual diffusion toward the roots (Figure **S1**). Plates were sealed with parafilm and mounted on a benchtop rotator (ELMI Intelli-Mixer™ RM-2L) rotating at 9 RPM to achieve continuous clinorotation during the assay. For chemical treatments, seedlings were placed on medium containing ascorbic acid (0.5 mM in DDW) or diphenyleneiodonium (DPI; 1 μM in DMSO) 1 h prior to halostimulation. Halostimulation was then initiated by adding the NaCl-containing half of the split-agar system to the medium. Seedlings were imaged using a Nikon D7100 camera with an AF-S DX Micro NIKKOR 85 mm f/3.5G ED VR lens (Nikon, Tokyo, Japan), and root-tip curvature was quantified using ImageJ software (https://imagej.nih.gov/ij/).

### Confocal Microscopy and ROS Detection

ROS accumulation in roots was visualized using dihydrorhodamine 123 (DHR) as previously described (Krieger *et al*. 2016). Briefly, immediately following halotropic stimulation, seedlings were immersed in 86.5 μM DHR (0.003% w/v; Sigma-Aldrich) prepared in phosphate-buffered saline (PBS, pH 7.4) for 2 or 5 min. After staining, seedlings were briefly rinsed in PBS to remove excess dye and mounted in PBS on glass slides. Confocal imaging was performed using a Leica SP8 laser-scanning confocal microscope (https://www.leica-microsystems.com/products/confocal-microscopes/p/leica-tcs-sp8) equipped with a 10× air objective. Imaging settings were kept constant across all samples to allow quantitative comparison. Excitation was at 488 nm (2% laser power), and emission was collected between 519 and 560 nm. Detector master gain was maintained between 670 and 720 (instrument-specific arbitrary units), with digital gain set to 1. Images were acquired using identical pinhole size, scan speed, and resolution parameters for all treatments. Fluorescence intensity was quantified as mean gray value using ImageJ/Fiji. For length measurements, ROS domains were defined manually using segmented region-of-interest tools, applying the same thresholding criteria across samples. Confocal images were pseudo-colored (RGB LUT) in ImageJ for visualization purposes only, without altering raw pixel values.

### Statistical Analysis and Data Presentation

Data were analyzed using OriginPro 2024 (OriginLab Corporation) and Microsoft Excel 2021 with the Analysis ToolPak add-in. Graphs and summary statistics were generated using these software packages, and all statistical tests are indicated in the figure legends or main text.

### Accession numbers

Accession numbers of the major genes investigated in this research are: *RBOHC,* AT5G51060; *GNOM*, AT1G13980.

## Results

### Analysis of Halotropic Bending under Minimized Gravitropic Influence

To isolate the halotropic response from gravitropic effects, wild-type seedlings were placed in the NaCl/split-agar system for halostimulation under continuous clinorotation (see Materials and Methods, and Figures **1a** and **S1a**). Briefly, seedlings were initially grown on control medium and then positioned at the interface between the control agar and the NaCl-containing agar, creating a defined directional salt cue, and root-bending angles were quantified under continuous clinorotation. This approach has been previously validated for minimization of the directional gravity vector in studies of tropic responses (Wexler *et al*. 2024; Zhang *et al*. 2025). Under normal gravity conditions, roots exposed to a lateral salt gradient exhibited a halotropic response accompanied by gravitropic growth (Figure **1a**). The NaCl/split-agar system was applied continuously under both normal gravity and uninterrupted clinorotation conditions (Figure **S1b**), enabling a direct comparison of bending dynamics between normal gravity and substantially reduced gravity conditions, while minimally perturbing the growth environment.

**Figure 1.**
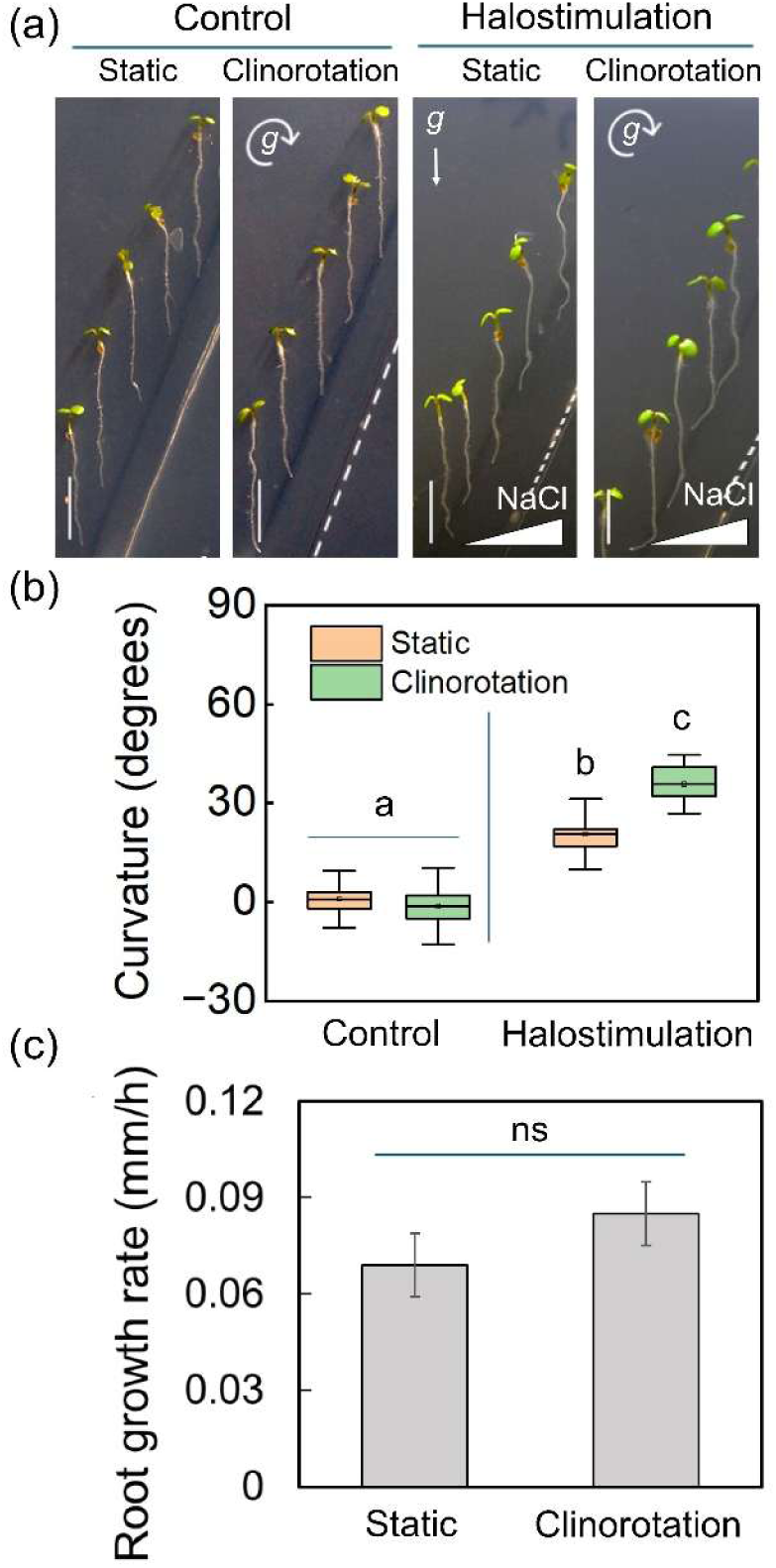
Clinorotation enhances halotropic bending in *Arabidopsis* roots. Seven-day-old wild-type *Arabidopsis* (Col-0) seedlings were halostimulated in a NaCl/split-agar system for 12 h under either static (control gravity) or continuous clinorotation conditions. (a) Representative images showing root-tip curvature. Scale bar = 0.5 cm; g, gravity vector, with direction indicated by arrow. (b) Quantification of root curvature. Box plots indicate the interquartile range; horizontal lines represent the median; whiskers denote the standard deviation (SD). Different letters indicate significant differences (Tukey’s HSD post hoc test, *P* < 0.001). (c) Root-elongation rate determined by measuring primary root length at the beginning and end of the halostimulation period. Values represent mean ± SD (three independent experiments of 15 seedlings each; *n* = 45). No significant differences (ns) were detected between treatments according to Student’s *t*-test.

Quantification of the bending angles (Figure **1b**) showed minimal curvature in the absence of salt (static control: 0.9 ± 4.3 degrees, mean ± SD). Salt stimulation significantly increased bending under static conditions (static halostimulation: 20.6 ± 5.4 degrees). When gravitropic input was minimized by continuous clinorotation, basal curvature remained close to zero (clinorotation control: –1.3 ± 5.8 degrees), whereas salt stimulation elicited a markedly stronger halotropic response (clinorotation halostimulation: 35.7 ± 4.4 degrees). Thus, removal of the gravitational vector substantially increased the magnitude of halotropic bending relative to static conditions.

Importantly, root-elongation rates measured during the assay (Figure **1c**) showed that clinorotation leads to slightly faster elongation compared to static growth, although this difference was not statistically significant. Therefore, the enhanced halotropic response under clinorotation cannot be attributed to altered growth rates but instead, likely reflects suppression of the competing gravitropic machinery.

Together, these findings indicated that clinorotation promotes halotropic curvature primarily by attenuating gravitropic signaling, thereby allowing the intrinsic dynamics and full magnitude of the halotropic response to emerge.

### NaCl/Split-Agar Halostimulation Induces a Spatially Restricted ROS Domain that Expands under Clinorotation

ROS have been shown to tune the balance between gravitropism and hydrotropism (Krieger *et al*., 2016). To assess whether analogous oxidative mechanisms contribute to halotropism, we analyzed ROS distribution in *Arabidopsis* roots exposed to a lateral salt gradient under continuous clinorotation.

ROS accumulation can be monitored using the H₂O₂-sensitive probe DHR (Gomes, Fernandes & Lima 2005), allowing visualization of dynamic, root-zone-specific ROS patterns (Royall & Ischiropoulos 1993; Crow 1997; Krieger *et al*. 2016). Confocal microscopy imaging of DHR-stained wild-type roots revealed a strong and symmetric ROS signal confined to the epidermal cells of the elongation zone (Figure **2a**). This symmetric oxidative domain was absent from the meristem and other tissues, making it a characteristic feature associated with directional salt-stimulated root growth. Quantification of fluorescence intensity (Figure **2b**) showed that asymmetric salt exposure markedly increases ROS accumulation (arbitrary units, mean ± SD). Under static conditions, values rose from 2170 ± 505.6 (control) to 4250 ± 510.4 following halostimulation, whereas under clinorotation, ROS levels increased from 2290 ± 443.3 (control) to 4310 ± 550.7. These data confirmed that formation of the oxidative domain depends on directional salt input rather than uniform exposure. We next quantified the longitudinal extent of the ROS-enriched region. In both static and clinorotated roots, the oxidative signal consistently initiated at approximately 400 μm from the root tip. However, the length of the ROS domain differed markedly between conditions: under static growth, it measured 293 ± 40 μm, whereas under clinorotation it expanded to 565 ± 67 μm, representing a significant increase (Figure **2c**). These measurements indicated that reducing gravitropic input promotes a substantially more extended oxidative domain within the elongation zone.

**Figure 2.**
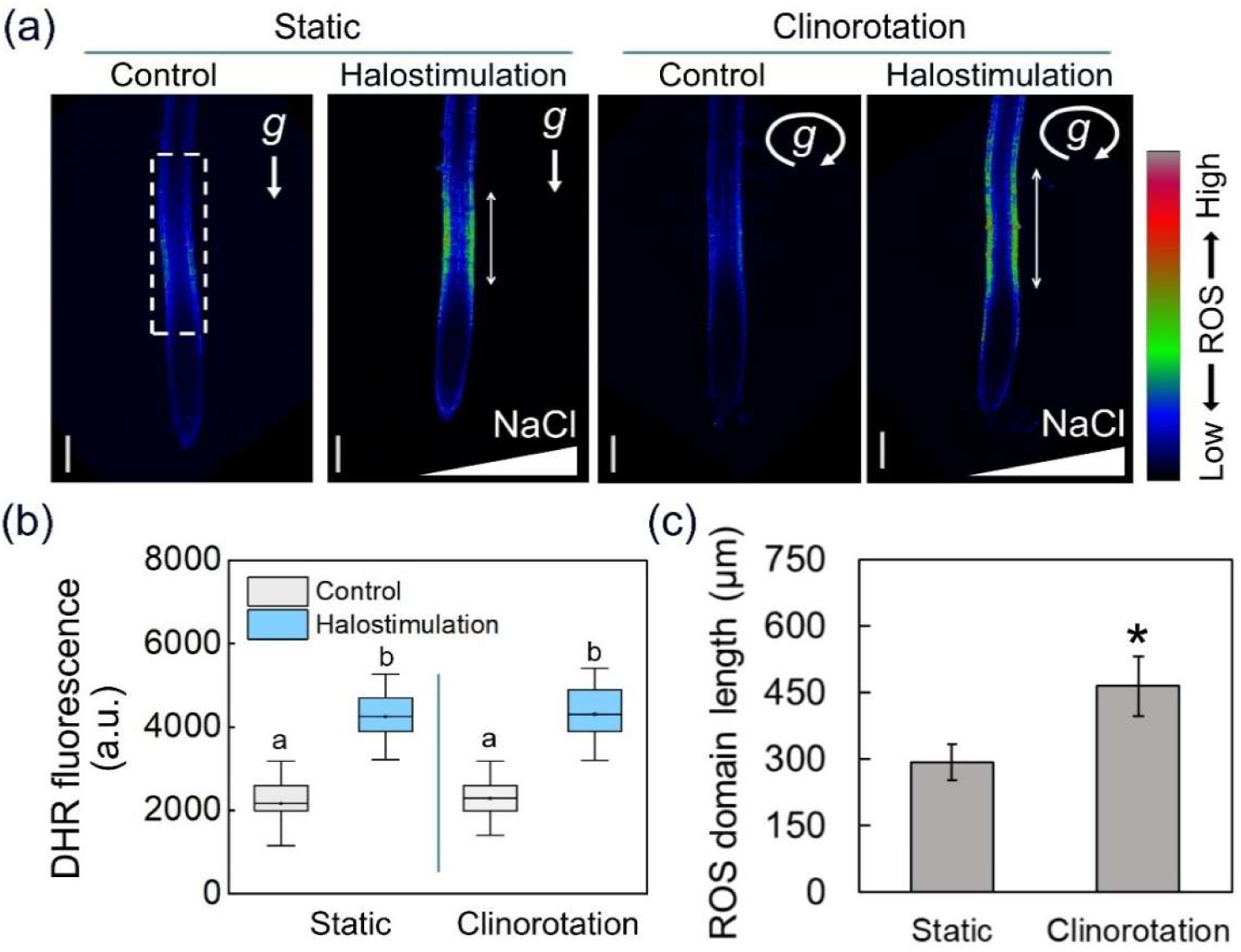
Exposure of *Arabidopsis* roots to a salt gradient evokes an elongation-zone-engulfing, gravity-responsive ROS signal. (a–c) Four-day-old *Arabidopsis* seedlings were either kept under control conditions or subjected to halostimulation for 90 min, under static vertical orientation or continuous clinorotation to minimize gravitropic effects. (a) Representative DHR-staining images, visualized by confocal microscopy and pseudo-colored to show relative ROS levels. Scale bar = 100 µm; g, gravity vector, with direction indicated by arrow. (b) Quantification of DHR fluorescence intensity in 300–900 µm root segments from the apex (dashed box in panel a; a.u., arbitrary units). Box plots show interquartile range; horizontal lines indicate the mean; whiskers denote SD. Different letters indicate significant differences (Tukey’s HSD post hoc test, *P* < 0.001). (c) Quantification of ROS-domain length as indicated by the double-headed arrow in panel a. Error bars show mean ± SD (three independent experiments, 10 seedlings each; *n* = 30). Asterisk indicates significant difference between treatments (Student’s *t*-test, *P* < 0.01).

Control experiments confirmed the specificity of the ROS response to directional salt exposure. When seedlings were subjected to a vertical salt gradient applied from below toward the root tips (Figure **S2a**), roots displayed only diffuse, non-localized bending (Figure **S2b**) and lacked the defined symmetric ROS domain (Figure **S2c**), with quantification showing no significant signal accumulation (Figure **S2d**). Similarly, exposure to homogeneous NaCl concentrations from all directions (schematized in Figure **S3a**) failed to induce localized ROS accumulation. Roots treated with 25 mM or 50 mM NaCl—mimicking the approximate concentrations reached in the elongation zone after 1–2 h of halostimulation (Figure **S1**)—showed no discernible ROS signal (Figure **S3b**) and quantification confirmed the absence of localized fluorescence (Figure **S3c**).

Together, these observations identified a symmetric, spatially restricted ROS domain in the elongation zone that forms specifically under directional, lateral salt exposure as provided by the NaCl/split-agar halostimulation system (Figure **S1**), and that cannot be reproduced by vertical gradients or uniform NaCl treatments (Figures **S2** and **S3**). These findings indicated that the formation of the ROS domain in the elongation zone specifically requires a side-directed halostimulus.

### RBOHC-Dependent ROS Modulate Halotropic Bending in *Arabidopsis* Roots

To assess the potential involvement of RBOHC, previously shown to mediate ROS production during hydrotropism (Krieger *et al*. 2016), we compared root halotropic responses and ROS accumulation in wild-type and *rbohC*-mutant seedlings halostimulated in the NaCl/split-agar system (Figure **3a**). Under halostimulation, *rbohC* roots exhibited a markedly exaggerated bending response, reaching 59.7 ± 6.3 degrees, significantly exceeding the 34.9 ± 8.1 degrees observed in the wild-type (Figure **3b**). In contrast, under control conditions, both genotypes showed minimal curvature, with wild-type roots bending 0.5 ± 5.8 degrees and *rbohC* roots 0.2 ± 5.5 degrees. Root-elongation rates during the assay were similar between genotypes (Figure **3c**), indicating that the enhanced curvature of *rbohC* under halostimulation is not attributable to differences in growth rate.

**Figure 3.**
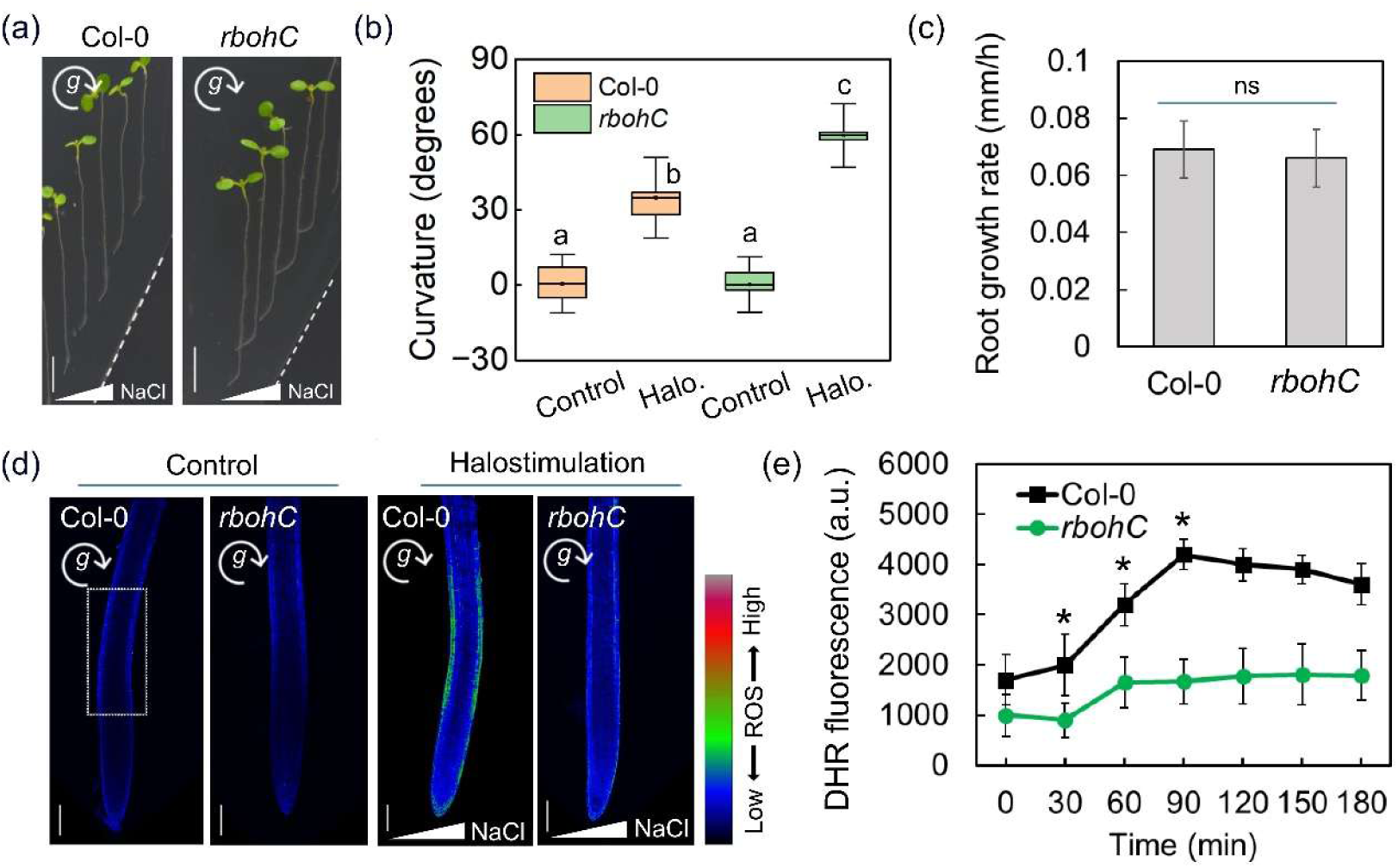
RBOHC is required for the formation of the halotropically induced ROS signal. (a–d) Five-day-old wild-type (Col-0) and *rbohC* mutant *Arabidopsis* seedlings were exposed to halostimulation (Halo.) in a NaCl/split-agar system for 12 h under continuous clinorotation. (a) Representative images of root-tip curvature. Scale bar = 0.5 cm; g, gravity vector, with direction indicated by arrow. (b) Quantification of root curvature. Box plots represent the interquartile range; horizontal lines indicate the mean; whiskers denote SD (three independent experiments of 15 seedlings each; *n* = 45). Different letters indicate significant differences by Tukey’s HSD (*P* < 0.001). (c) Root-elongation rate calculated from primary root length at the start and end of halostimulation. Values represent mean ± SD (three independent experiments of 15 seedlings each; *n* = 45); ns, no significant difference relative to Col-0 (Student’s *t*-test). (d) Representative DHR-staining images of Col-0 and *rbohC* roots under control and halostimulated conditions after 90 min, visualized by confocal microscopy and pseudo-colored to indicate relative ROS levels. Scale bar = 100 µm. (e) Quantification of DHR fluorescence intensity in the elongation zone at the indicated time points (dashed box in panel d; a.u., arbitrary units). The line plot shows mean values with whiskers indicating SD. Asterisks indicate significant differences between Col-0 and the mutant seedlings (Student’s *t*-test, *P* < 0.001).

Consistent with these phenotypes, DHR staining revealed a strong ROS signal in the elongation zone of wild-type roots, peaking approximately 90 min after halostimulation (Figure **3d,e**). In *rbohC* seedlings, no comparable ROS accumulation was detected during this time course, with basal fluorescence remaining low. Quantification confirmed that ROS intensity in wild-type roots was roughly fourfold higher than in the mutant. These findings indicated that RBOHC is required for generation of the halotropically induced ROS signal. Notably, the absence of ROS in *rbohC* correlated with an exaggerated halotropic curvature, suggesting that ROS serve as a regulatory mechanism modulating the magnitude of the root-bending response and preventing excessive curvature in response to lateral salt stimuli.

In contrast, *rbohD* seedlings did not exhibit any significant differences in halotropic curvature or root elongation compared to the wild-type (Figure **S4**). RBOHD is expressed predominantly in stems and leaves and functions primarily in systemic ROS signaling (Sagi & Fluhr 2006; Miller *et al*. 2009; Suzuki *et al*. 2011). This is consistent with previous observations in hydrotropism, where RBOHC—but not RBOHD—was specifically required for ROS-mediated root responses (Krieger et al., 2016), highlighting a conserved and root-specific role for RBOHC in tropic signaling.

### Chemical Reduction of ROS Enhances Root Halotropism

To further investigate the role of ROS in root halotropism, we evaluated the effects of the ROS scavenger ascorbate and the NADPH oxidase inhibitor DPI, both known to suppress ROS accumulation in roots and previously shown to inhibit gravitropism while enhancing hydrotropism (Joo, Bae & Lee 2001; Foreman *et al*. 2003; Peer, Cheng & Murphy 2013; Krieger *et al*. 2016). Seedlings were subjected to halostimulation in the NaCl/split-agar assay with the medium segment carrying the roots containing either 0.5 mM ascorbate or 1 μM DPI. Both chemical treatments markedly enhanced root halotropism in wild-type seedlings compared to control-treated seedlings (Figure **4a,b**). Under halostimulation in the NaCl/split-agar system, DPI increased the bending angle to 51.3 ± 7.2 degrees, and ascorbate produced a similar enhancement, reaching 54.6 ± 8.2 degrees. In contrast, control-treated roots bent only 30.2 ± 6.3 degrees under the same conditions. These findings demonstrated that reducing ROS levels potentiates the halotropic response, supporting a model in which ROS act as modulators that constrain the magnitude of root bending during halotropism. Importantly, this enhancement of halotropic bending occurred despite a significant reduction in root-elongation rates by both chemicals (Figure **4c**), indicating that the increased curvature does not arise from accelerated growth but from modulation of the signaling machinery underlying the response.

**Figure 4.**
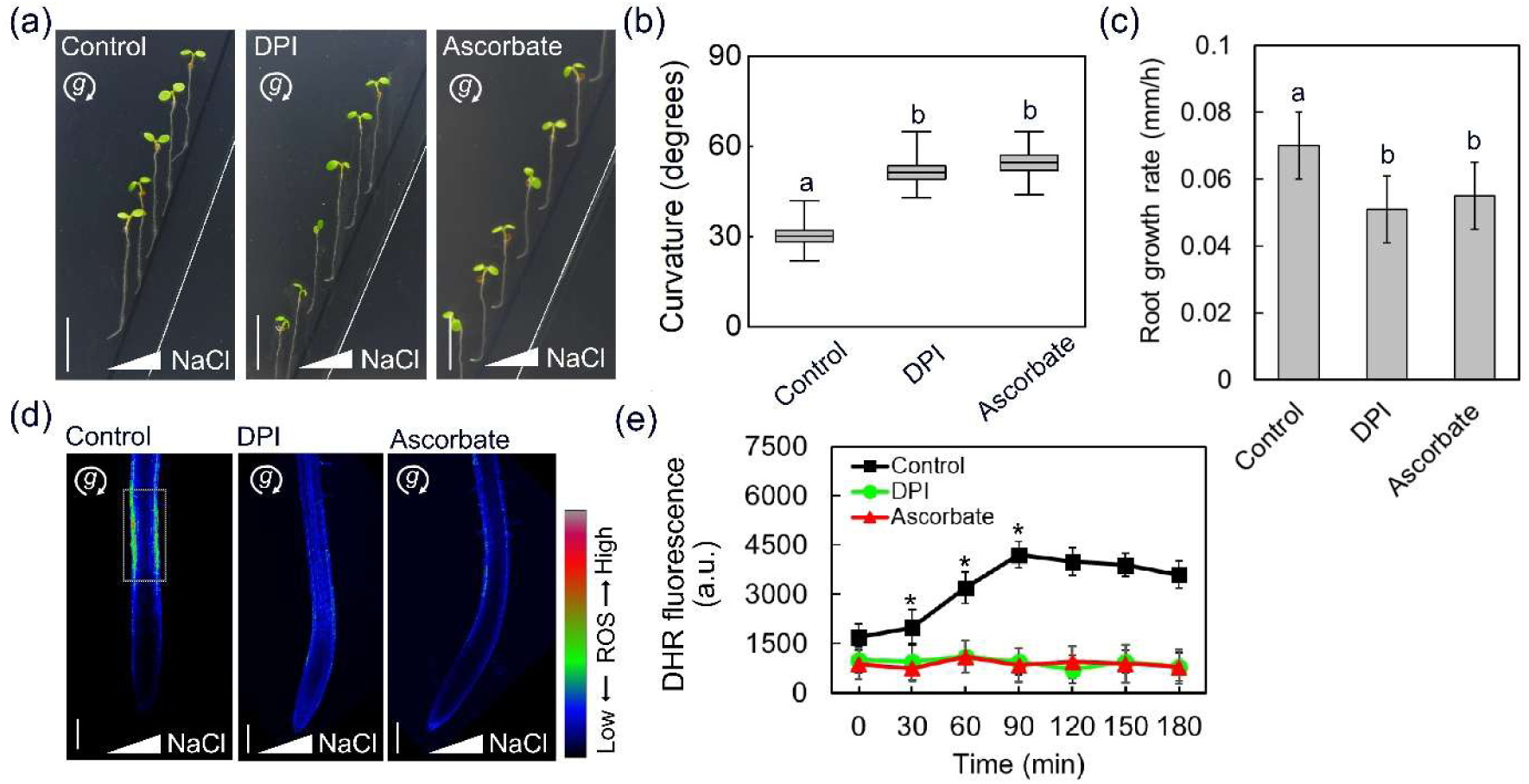
Reducing ROS levels enhances root halotropism. (a) Five-day-old wild-type (Col-0) *Arabidopsis* seedlings were exposed to a lateral salt gradient in a NaCl/split-agar system for 12 h under control conditions or in the presence of 0.5 mM ascorbate or 1 μM DPI. Representative images show root-tip curvature. Scale bar = 1 cm; g, gravity vector, with direction indicated by arrow. (b) Quantification of root-tip curvature for seedlings treated as in panel a. Box plots represent the interquartile range; horizontal lines indicate the mean; whiskers denote SD. Significant differences were determined by Tukey’s HSD (*P* < 0.01). (c) Root-elongation rate measured at the beginning and end of halostimulation. Values represent mean ± SD (three independent experiments of 15 seedlings each; *n* = 45); ns, no significant difference relative to Col-0 (Student’s *t*-test). (d) Representative DHR-staining images of halostimulated Col-0 roots under control, ascorbate, or DPI treatments after 90 min of clinorotation. Images were acquired by confocal microscopy and pseudo-colored to reflect relative ROS levels. Scale bar = 100 µm. (e) Quantification of DHR fluorescence intensity in the elongation zone at the indicated time points (dashed box in panel d; a.u., arbitrary units). The line plot shows mean values with whiskers indicating SD. Asterisks indicate significant differences relative to control (Student’s *t*-test, *P* < 0.001).

DHR staining of halostimulated seedlings revealed a clear correspondence between chemical suppression of ROS and absence of the oxidative domain in the elongation zone. Under continuous clinorotation in the NaCl/split-agar system, control-treated roots displayed the characteristic ROS-enriched region, whereas this domain was completely abolished in DPI-and ascorbate-treated seedlings (Figure **4d**). Fluorescence quantification along the elongation zone corroborated this pattern: control roots exhibited the typical transient rise in ROS, peaking at ∼90 min and remaining relatively stable through 180 min, whereas both treatments strongly attenuated ROS accumulation across the entire time course (Figure **4e**). These results mirror those obtained with the *rbohC* mutant and reinforce the conclusion that the halotropically induced ROS signal acts as a regulatory mechanism limiting the intensity of the halotropic response.

### *miz2* Mutants Exhibit Negative Halotropism and Aberrant ROS Distribution

Hydrotropism-associated genes are known to shape root responses to asymmetric water availability, and *miz1* mutant maintains a normal halotropic response (Yu *et al*. 2022). We therefore examined how the hydrotropism-defective mutant *miz2* responds to a lateral salt cue. In the NaCl/split-agar assay, *miz2* roots exhibited a pronounced reversal of growth direction: instead of bending away from the saline medium, they consistently grew toward it, displaying clear negative halotropism. After 16 h of halostimulation under continuous clinorotation, wild-type roots showed a positive curvature of 40.3 ± 7.8 degrees, whereas *miz2* roots bent toward the salt source, reaching −11.8 ± 5.7 degrees (Figure **5a,b**). In the corresponding no-salt control, wild-type roots displayed negligible curvature (0.11 ± 4.9 degrees) and *miz2* roots showed a similarly minimal response (0.22 ± 1.2 degrees), indicating that the negative curvature arises specifically under halostimulation. Root-elongation rates did not differ significantly between genotypes (Figure **5c**), demonstrating that the reversed bending reflects altered directional growth rather than differences in overall growth capacity.

**Figure 5.**
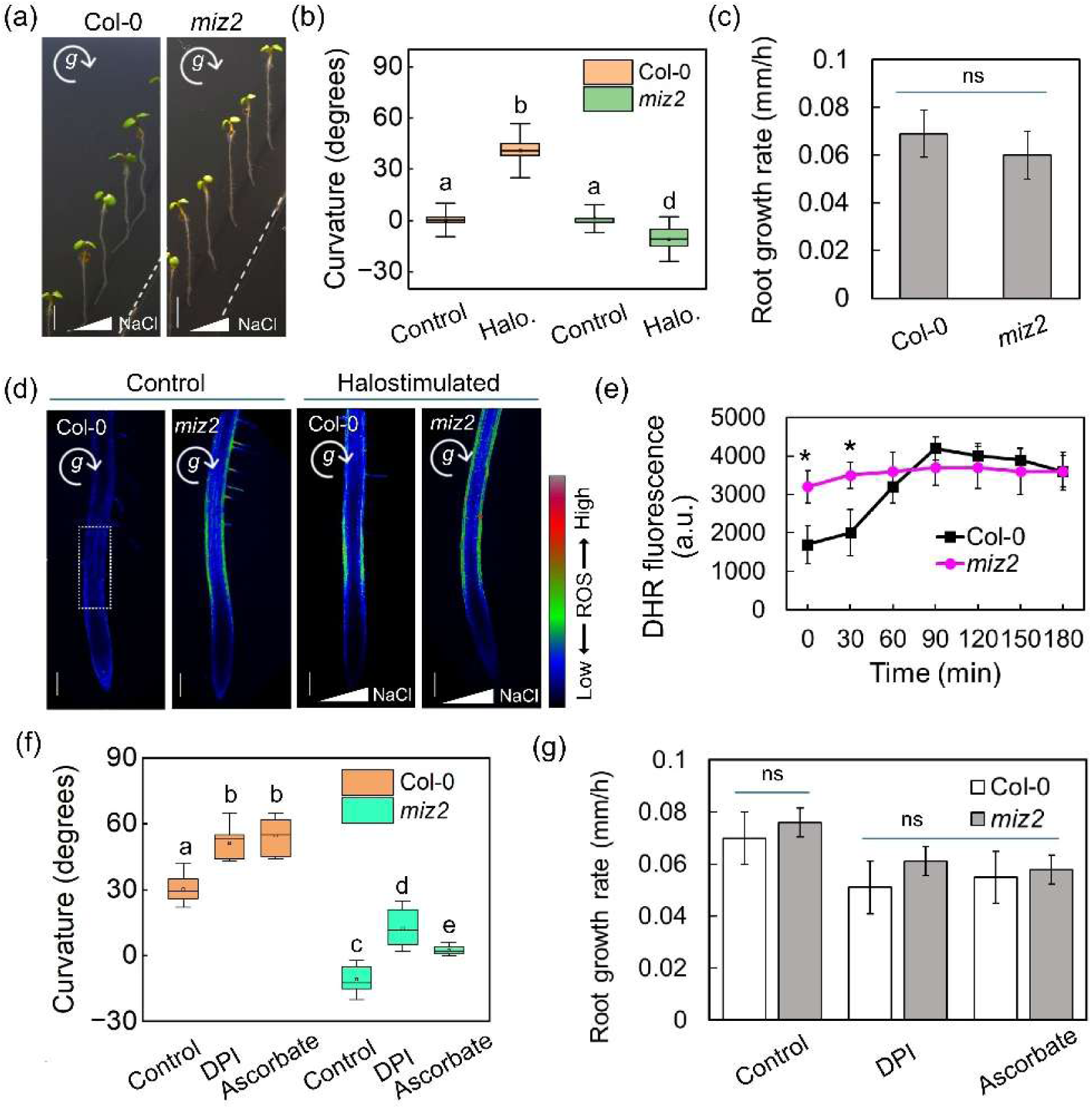
Negative halotropism and altered ROS distribution in *miz2* mutant roots. (a) Five-day-old wild-type (Col-0) and *miz2* mutant *Arabidopsis* seedlings were exposed to a lateral NaCl gradient in a NaCl/split-agar system for 16 h under continuous clinorotation. Scale bar = 0.5 cm; g, gravity vector, with direction indicated by arrow. (b) Quantification of root-tip curvature for halostimulated seedlings shown in panel a. Box plots represent the interquartile range; horizontal lines indicate the mean; whiskers denote SD. Significant differences were determined by Tukey’s HSD (*P* < 0.001). (c) Root-elongation rate measured at the beginning and end of halostimulation. Values represent mean ± SD (three independent experiments of 15 seedlings each; *n* = 45); ns, no significant difference relative to Col-0 (Student’s *t*-test). (d) Representative DHR-staining images of Col-0 and *miz2* roots under control and halostimulated conditions. Images were acquired by confocal microscopy and pseudo-colored to visualize ROS accumulation. Scale bar = 100 µm. (e) Quantification of DHR fluorescence intensity in the elongation zone (dashed box in panel d; a.u., arbitrary units). The line plot shows mean values with whiskers indicating SD (three independent experiments of 10 seedlings each; *n* = 30). Asterisks indicate significant differences relative to Col-0 (Student’s *t*-test, *P* < 0.001). (f) Quantification of root-tip curvature in halostimulated seedlings treated with control solution, ascorbate (0.5 mM), or DPI (1 µM). Box plots are presented as in panel b. Significant differences were determined by Tukey’s HSD (*P* < 0.001). (g) Root-elongation rate measured as in panel c for seedlings subjected to control, ascorbate, or DPI treatments. Values represent mean ± SD (three independent experiments of 15 seedlings each; *n* = 45); ns, no significant difference relative to Col-0 (Student’s *t*-test).

To determine whether this phenotype is associated with altered ROS distribution, we analyzed DHR fluorescence in halostimulated roots. In *miz2* seedlings, ROS accumulated under control conditions across a broad ∼400–1000 µm region from the apex. Following halostimulation, the ROS signal extended from ∼400 µm from the apex throughout the entire root segment captured within the imaging frame (Figure **5d,e**). In contrast, wild-type roots exhibited a halotropically induced ROS domain restricted to a defined ∼400–700 µm region of the elongation zone. This aberrant ROS pattern in *miz2* provides a physiological correlate for its reversed bending behavior and reflects altered spatial ROS dynamics during halotropism.

Next, we tested whether chemically lowering ROS levels would modulate this atypical response. In wild-type roots, DPI and ascorbate markedly enhanced halotropic bending, increasing curvature from 30.2 ± 6.3 degrees in control-treated seedlings to 51.3 ± 7.2 and 54.6 ± 8.2 degrees, respectively. In *miz2* roots, which exhibited negative bending under control conditions (−10.8 ± 6.5 degrees), DPI partially restored positive curvature to 12.4 ± 8.2 degrees, whereas ascorbate produced a smaller effect (2.4 ± 2.1 degrees) (Figure **5f**). Analysis of root-elongation rates across both genotypes and treatments revealed a similar reduction in growth for wild-type and *miz2* in the presence of either chemical, indicating that the observed differences in halotropic curvature are not due to altered growth rates (Figure **5g**). These results further indicated that the abnormal ROS distribution in *miz2* contributes to its atypical root bending under halostimulation.

### Loss of RBOHC Suppresses the Negative Halotropism and Aberrant ROS Pattern of *miz2*

To determine whether the altered halotropic response of *miz2* depends on RBOHC-mediated ROS production, we generated a *miz2 rbohC* double mutant by crossing the two single mutants, and analyzed its behavior under halostimulation. When halostimulated in the NaCl/split-agar assay, wild-type roots displayed a strong positive curvature away from the salt source (42.8 ± 7.8 degrees), whereas *miz2 rbohC* roots showed a markedly attenuated response of 10.4 ± 7.1 degrees (mean ± SD; Figure **6a,b**). Although the magnitude of the curvature in the double mutant was reduced relative to the wild-type, its bending was consistently positive, in sharp contrast to the negative curvature of *miz2* (Figure **5a,b**) and the enhanced curvature of *rbohC* (Figure **3a,b**). Thus, eliminating RBOHC activity genetically mitigated the directional defect of *miz2*.

**Figure 6.**
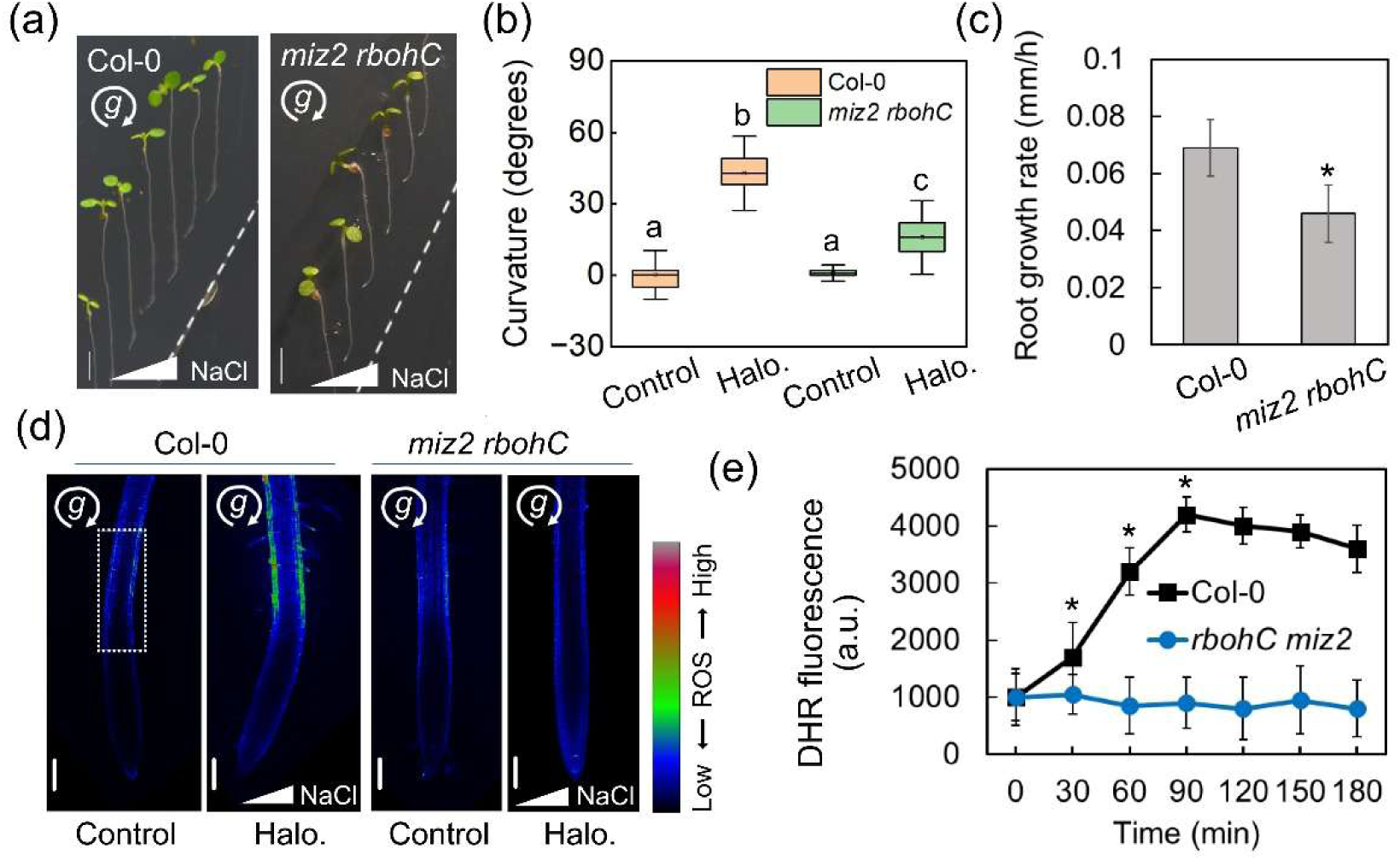
Partial restoration of halotropism in the *miz2 rbohC* double mutant. (a) Five-day-old wild-type (Col-0) and *miz2 rbohC* mutant *Arabidopsis* seedlings were exposed to a lateral NaCl gradient in a NaCl/split-agar system for 16 h under continuous clinorotation. Scale bar = 0.5 cm; g, gravity vector, with direction indicated by arrow. (b) Quantification of root-tip curvature for halostimulated (Halo.) seedlings shown in panel a. Box plots represent the interquartile range; horizontal lines indicate the mean; whiskers denote SD. Significant differences were determined by Tukey’s HSD (*P* < 0.001). (c) Root-elongation rate measured at the beginning and end of halostimulation. Values represent mean ± SD (three independent experiments of 10 seedlings each; *n* = 30). Asterisks indicate significant differences relative to Col-0 (Student’s *t*-test, *P* < 0.01). (d) Representative DHR-staining images of Col-0 and mutant roots under control and halostimulated conditions. Images were acquired by confocal microscopy and pseudo-colored to visualize ROS accumulation. Scale bar = 100 µm; g, gravity vector, with direction indicated by arrow. (e) Quantification of DHR fluorescence intensity in the elongation zone (dashed box in panel d; a.u., arbitrary units). Values represent mean ± SD (three independent experiments of 10 seedlings each; *n* = 30). Asterisks indicate significant differences relative to the corresponding Col-0 value at each time point (Student’s *t*-test, *P* < 0.001).

Under control (no-salt) conditions, both genotypes showed minimal curvature (1.2 ± 6.0 degrees in the wild-type; 0.9 ± 1.7 degrees in the double mutant), confirming that the restored positive bending of *miz2 rbohC* is specifically elicited by halostimulation. Analysis of root elongation revealed that *miz2 rbohC* roots elongate significantly more slowly than wild-type roots (Figure **6c**), indicating that their partially restored directional response does not arise from increased growth capacity.

To assess the underlying ROS pattern, we examined DHR fluorescence. Unlike *miz2*, which exhibited a broad and uncontrolled ROS domain, the *miz2 rbohC* double mutant showed no detectable ROS accumulation under halostimulation (Figure **6d,e**). This complete loss of ROS signal provides strong evidence for generation of the aberrant ROS domain observed in *miz2* by RBOHC-dependent ROS production. Together, these results demonstrated that removing RBOHC-derived ROS suppresses both the mislocalized ROS accumulation and the negative halotropic curvature characteristic of *miz2*.

## Discussion

Our results indicate that halotropism involves a spatially restricted domain of RBOHC-generated ROS in the epidermis of the elongation zone, which forms during the first hours of lateral salt exposure and peaks in the primary root segment located approximately 400–700 µm from the root tip under normal gravity conditions (Figure **2**). Reducing ROS levels either chemically (DPI, ascorbate) (Figure **4**) or genetically (RBOHC knockout) enhanced halotropic curvature in wild-type roots (Figure **3**), in line with previous observations in hydrotropism (Krieger *et al*., 2016), suggesting that RBOHC-derived ROS act as a regulatory brake to prevent excessive bending. Importantly, our control experiments revealed that this ROS domain forms only in response to the side-specific halostimulation provided by the NaCl/split-agar system and is not induced by vertical gradients or uniform salt exposure (Figures **S2** and **S3**). Together, these findings indicate that the spatially restricted ROS signal fine-tunes directional growth by modulating gravitropic input, facilitating bending away from high-salinity regions, and acting as a modulatory rather than primary driver of halotropic curvature.

Clinorotation minimized gravitropic opposition and revealed the full amplitude of the halotropic bending (Figure **1**), while producing an appreciably extended ROS domain relative to normal gravity (Figure **2**). This expansion was sufficient to destabilize the ROS-based mechanism that normally constrains halotropic curvature, resulting in exaggerated bending (Figure **1a**). The pronounced enhancement of halotropic bending in response to the moderate expansion of the ROS domain indicates that ROS act as key regulators of curvature magnitude, and suggests that additional factors—potentially influenced by the reduction of gravitropic input—cannot be excluded. By reducing the effective gravitropic vector, clinorotation allows the halotropic cue to act with minimal antagonism from gravity. A reduction in ROS production—either chemically or genetically—similarly enhances halotropic curvature, consistent with roles for ROS in both curvature restraint and the maintenance of tropic responsiveness, as previously demonstrated for hydrotropism (Krieger *et al*. 2016). In *miz2*, the disruption of GNOM-dependent confinement produced a much more extensive and mislocalized ROS distribution, which not only removed ROS-dependent restraint but also prevented proper execution of halotropism. While clinorotation or ROS reduction primarily removed the ROS-dependent constraints on halotropic bending, the extensive mislocalization of ROS in *miz2* produced negative halotropism, highlighting the notion that ROS are not just passive elements, but function as active signaling components that modulate both the magnitude and direction of the halotropic response.

The *miz2* mutant, defective in ARF-GEF GNOM (Miyazawa *et al*. 2009), exhibited negative halotropism—bending toward the salt—together with a broad, constitutive, symmetric ROS domain extending across the elongation zone and the region above it (Figure **5a–e**). GNOM is a central regulator of vesicular trafficking and endosomal recycling (Geldner et al. 2003; Tanaka et al., 2009) and its disruption is suggested to affect the spatial distribution of RBOHC. Consistent with this idea, perturbation of vesicle trafficking using brefeldin A (BFA) has been shown to cause RBOHC accumulation in FM4-64-positive endocytic compartments and its removal from the plasma membrane, demonstrating the requirement for continuous endocytic turnover to maintain the spatial restriction of RBOHC to the cell surface (Takeda *et al*. 2008). In *miz2*, the broad, symmetric, and elongated ROS field may reflect altered distribution of RBOHC, preventing the spatial modulation of elongation required for halotropic bending away from salt. Together, these observations suggest that the restricted, symmetric ROS domain characteristic of wild-type roots is essential for proper halotropic curvature and for preventing excessive bending.

Chemical reduction of ROS partly restored positive halotropism in *miz2*, with DPI being more effective than ascorbate (Figure **5f,g**). The stronger rescue by DPI, which inhibits ROS production primarily via RBOH enzymes—including RBOHC, supports the notion that the *miz2* phenotype may arise from misregulated RBOHC activity or localization. Although ascorbate did not fully reconstitute the wild-type response, it effectively counteracted the negative bending (Figure **5f**), indicating that aberrant ROS accumulation is a major—but not exclusive—driver of the phenotype. Complete suppression of the expanded ROS domain and restoration of positive curvature were achieved by removing RBOHC in the *miz2* background (Figure **6**). The attenuated bending in *miz2 rbohC*, despite reduced growth rates (Figure **6c**), confirmed that the phenotype reflects altered directional growth rather than growth capacity.

Among the RBOH isoforms, RBOHC—but not RBOHD—emerges as essential for the generation of ROS in root-hair and elongation-zone cells. Consistent with its established role in root-hair ROS production (Foreman *et al*. 2003; Monshausen *et al*. 2007), RBOHC was required for the formation of the halotropically induced, elongation-zone ROS domain, as *rbohC* mutants lacked this localized signal and displayed exaggerated bending in response to a lateral salt gradient (Figure **3**). In contrast, *rbohD* mutants were indistinguishable from the wild-type (Figure **S4**), reflecting RBOHD’s predominant function in systemic ROS signaling (Sagi & Fluhr 2006; Miller *et al*. 2009; Suzuki *et al*. 2011). These observations highlight a conserved, local role for RBOHC in modulating tropic responses through spatially restricted ROS accumulation, extending its functional relevance from root-hair development to directional root growth under halostimulation.

Together, these results support a model in which GNOM-dependent trafficking confines RBOHC to defined plasma-membrane regions, generating a spatially restricted, symmetric ROS domain in the elongation zone that modulates the balance between tropic cues. In gravitropism, ROS accumulate asymmetrically, with higher levels on the lower side of the root to drive downward bending (Joo *et al*. 2001). In contrast, both hydrotropism and halotropism generate a symmetric, spatially defined ROS domain within the elongation zone (Krieger *et al*. 2016; Figure **2**), which suppresses gravitropic dominance and enables reorientation away from adverse conditions. Notably, although *miz1* is required for hydrotropism (Kobayashi *et al*. 2007) but not for halotropism (Yu *et al*. 2022), the formation of this defined ROS domain in both responses—and its disruption in *miz2*—suggest that GNOM functions upstream of a shared ROS-based signaling module that is essential for both tropisms.

Mechanical stimulation induces rapid and transient increases in cytosolic Ca²⁺, which act as the primary signal linking mechanical cues to downstream responses. These Ca²⁺ transients activate RBOHC, thereby triggering apoplastic ROS production at the cell wall with kinetics tightly coupled to the Ca²⁺ signal (Monshausen et al., 2009). Given the reported role of Ca²⁺ in hydrotropic signaling (Shkolnik *et al*. 2018), it is plausible that Ca²⁺ also contributes to the upstream regulation of RBOHC during halotropism, coordinating ROS-mediated modulation of root bending. Under clinorotation, this symmetric ROS domain becomes only modestly extended, yet this small expansion is sufficient to further weaken the gravitropic influence and reveal stronger halotropic curvature. In *miz2*, impaired spatial confinement leads to mislocalized ROS accumulation, shifting the response toward growth into the salt (Figure **5**). In contrast, *rbohC* shows the opposite behavior, with loss of RBOHC-derived ROS removing the restraint on curvature and causing exaggerated bending away from salt (Figure **3**). While our findings establish the requirement for GNOM-dependent spatial control of RBOHC activity, future approaches combining high-resolution imaging with targeted perturbation of vesicle trafficking will be required to refine the mechanistic model and directly assess trafficking-dependent localization dynamics.

Understanding the molecular logic underlying halotropic signaling is not only fundamental biologically, but also essential for developing crops capable of maintaining productivity in increasingly salt-affected agricultural soils—a growing challenge for sustainable agriculture and global food security.

## Acknowledgments

This research was supported by the Israel Science Foundation (grant no. 756/20 to D.S.).

## Author contributions

D.S., A.C. and M.F. designed the experiments and research plan; all authors performed the experiments, analyzed the data, and wrote the article.

## Data availability

The data that support the findings of this study are available from the corresponding author upon reasonable request.References

## Supplemental material

### Supporting information

**Figure S1.**
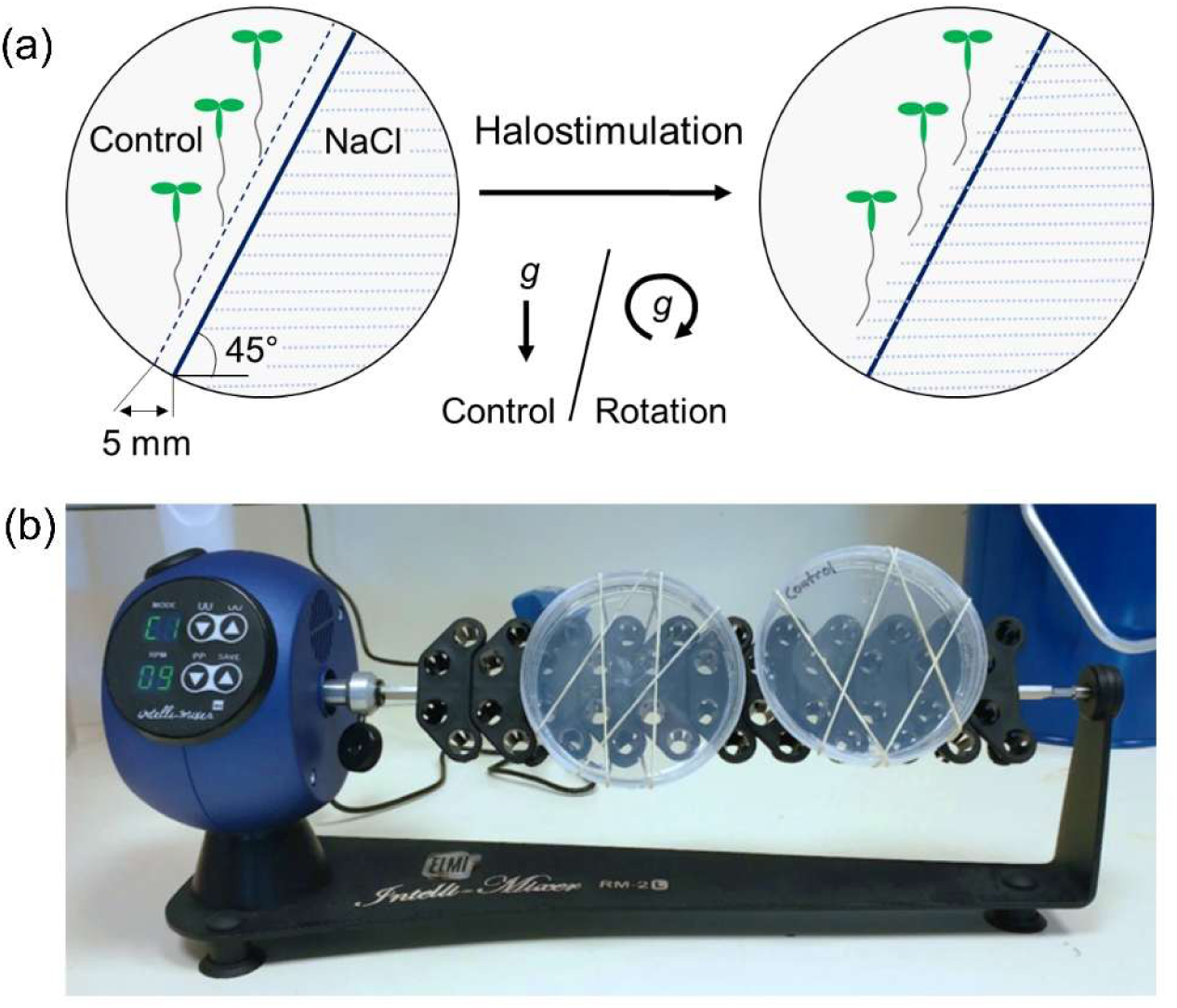
NaCl/split-agar system utilized for root halostimulation under control gravity and continuous clinorotation conditions. (a) Schematic of the NaCl/split-agar assay. Seedlings were placed diagonally on agar solidified with 0.25× MS medium occupying half of the plate area, followed by addition of 200 mM NaCl-containing medium to the other half. Root curvature was measured at specified time points after treatment initiation; g, gravity vector, with direction indicated by arrow. (b) Illustration of the rotation setup for clinorotation experiments. Assay plates were mounted on a rotator to minimize gravitropic influence during halostimulation.

**Figure S2.**
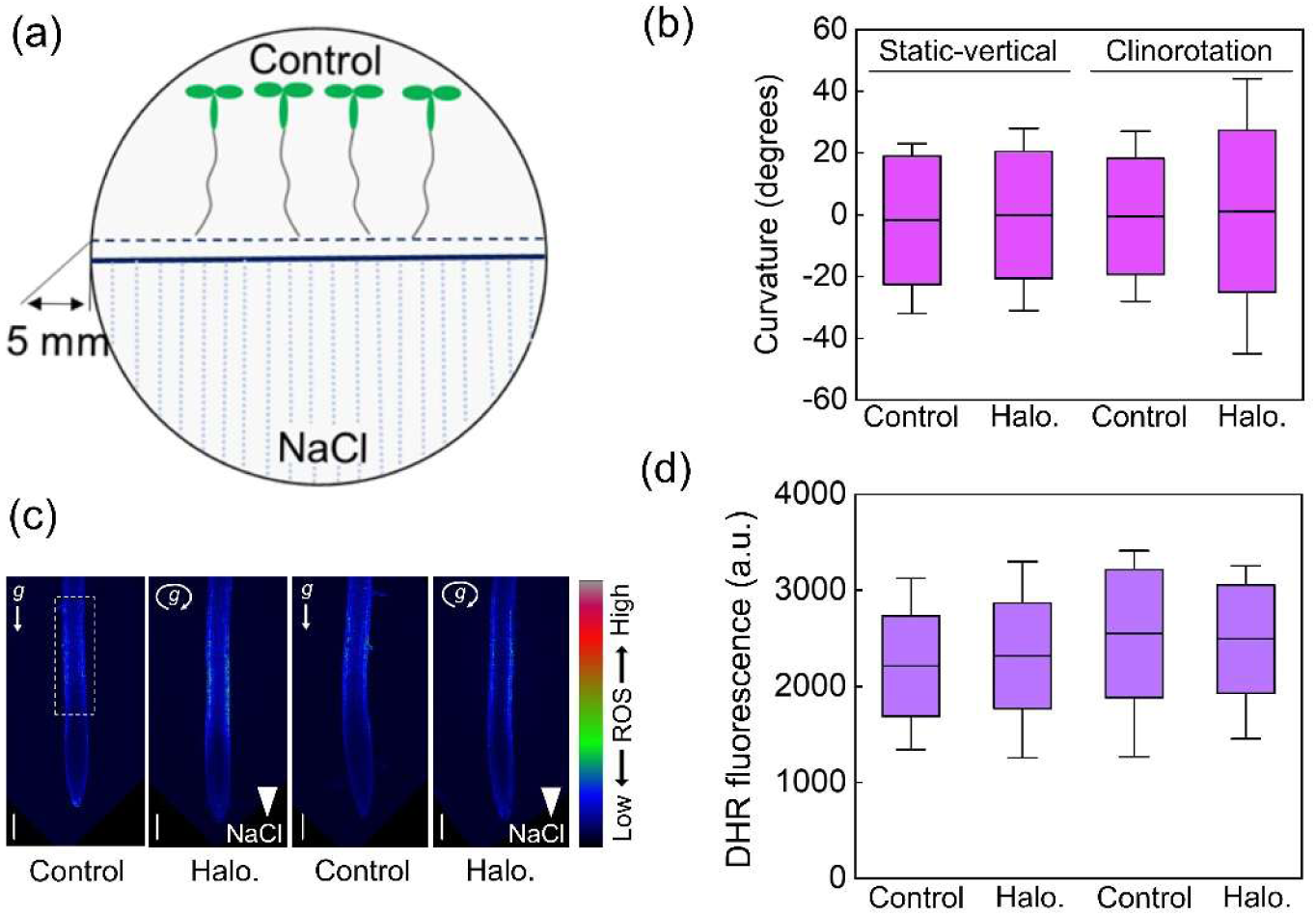
ROS signals are not generated in roots in response to horizontal halostimulation. (a) Schematic representation of the horizontal NaCl/split-agar system. Control plates were prepared by attaching half-plate medium without NaCl. (b) Quantification of root-tip curvature of 5-day-old wild-type (Col-0) *Arabidopsis* seedlings halostimulated (Halo.) in a static vertical position as depicted in panel a or under continuous clinorotation. Data represent mean ± SD of three independent experiments with 15 seedlings each (*n* = 15 total). (c) Representative DHR-staining images of roots from control and halostimulated seedlings. Images were pseudo-colored to indicate relative ROS levels. Scale bar = 100 µm; g, gravity vector, with direction indicated by arrow. (d) Quantification of DHR fluorescence intensity in 300–900 µm segments from the root apex (dashed box in panel c; a.u., arbitrary units). Data represent mean ± SD of three independent experiments with five seedlings each (*n* = 15 total).

**Figure S3.**
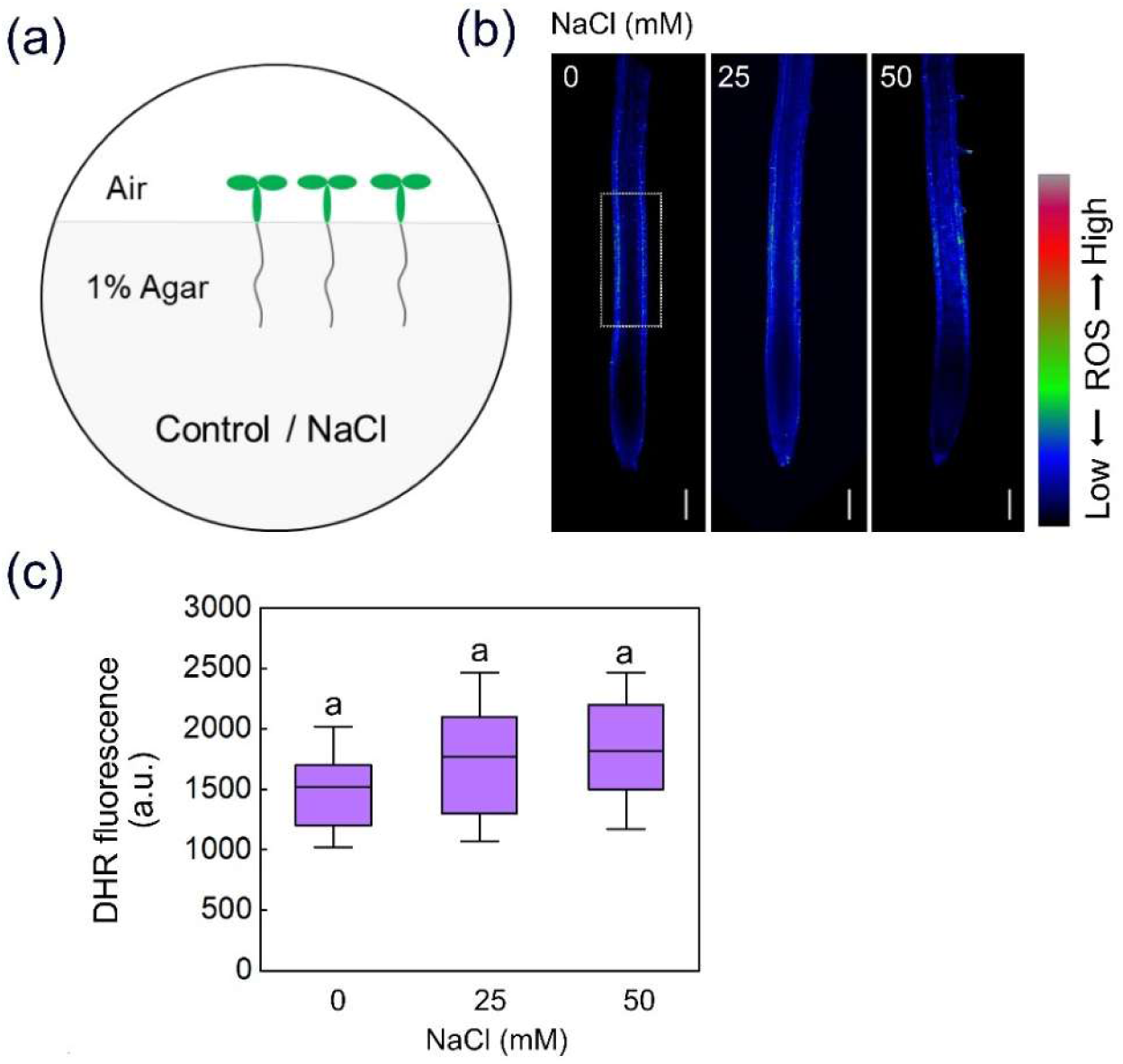
Uniform salt exposure does not induce ROS signal in the root-elongation zone. (a) Schematic representation of the experimental setup used to apply uniform salt stress to entire roots. (b) Representative staining of roots of 5-day-old wild-type (Col-0) *Arabidopsis* seedlings exposed to 0, 25, or 50 mM NaCl for 30 min. The region used for fluorescence quantification is indicated by a dashed box. (c) Quantification of DHR fluorescence intensity in the elongation zone (dashed box in panel b; a.u., arbitrary units). Bars represent mean ± SD (*n* = 30 roots). Uniform salt exposure did not induce ROS accumulation in this region, in contrast to the localized signal observed under halostimulation.

**Figure S4.**
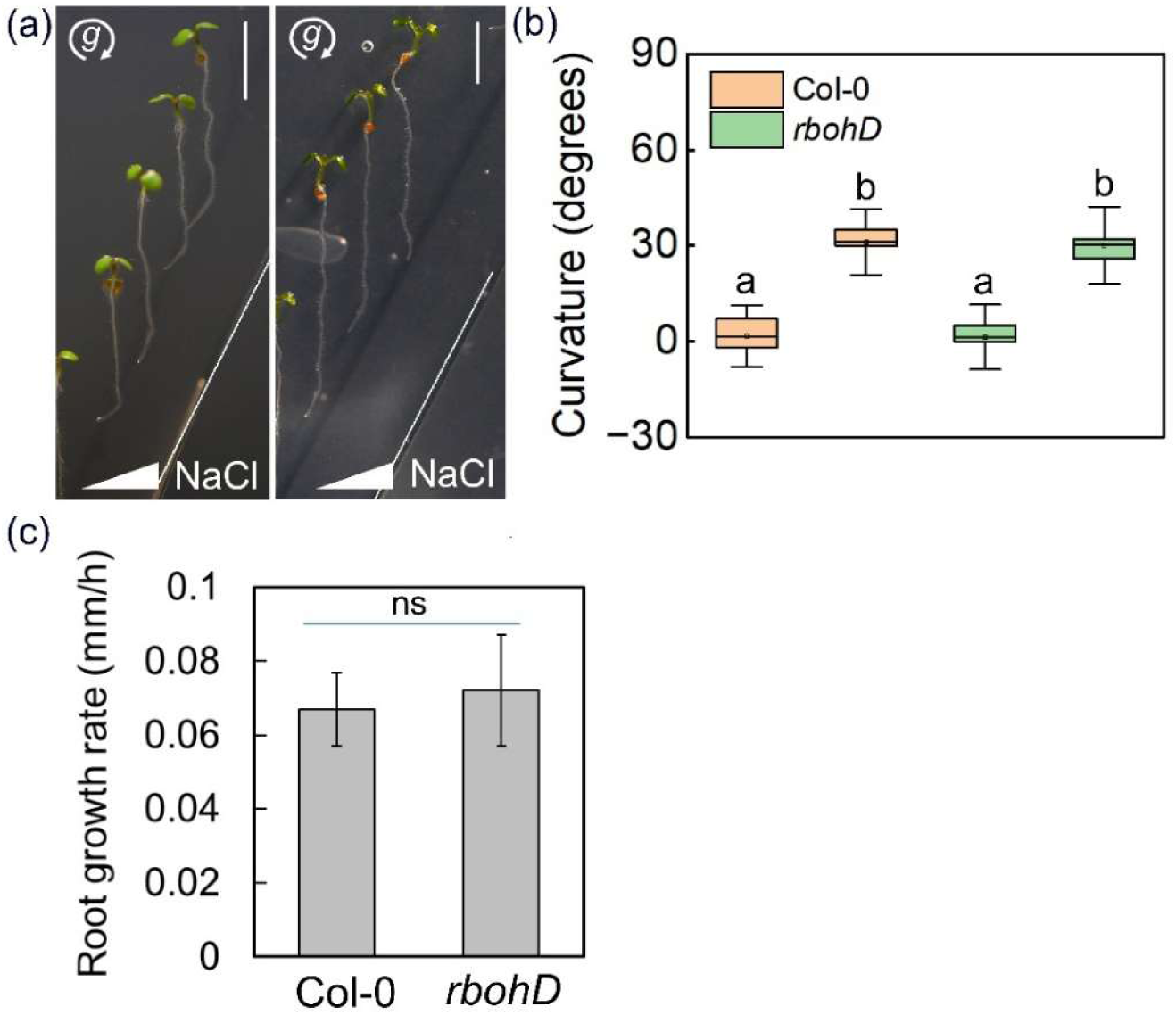
RBOHD is not required for the halotropically induced bending response. (a–d) Seven-day-old wild-type (Col-0) and *rbohD* mutant Arabidopsis seedlings were exposed to halostimulation (Halo.) in a NaCl/split-agar system for 12 h under continuous clinorotation. (a) Representative images of root-tip curvature. Scale bar = 0.5 cm; g, gravity vector, with direction indicated by arrow. (b) Quantification of root curvature. Box plots represent the interquartile range; horizontal lines indicate the mean; whiskers denote SD (three independent experiments of 15 seedlings each; *n* = 45). No significant differences were detected between Col-0 and *rbohD* (Tukey’s HSD, *P* > 0.05). (c) Root-elongation rate calculated from primary root length at the start and end of halostimulation. Values represent mean ± SD (three independent experiments of 15 seedlings each; *n* = 45). ns, no significant difference relative to Col-0 (Student’s *t*-test).

**Figure S5.**
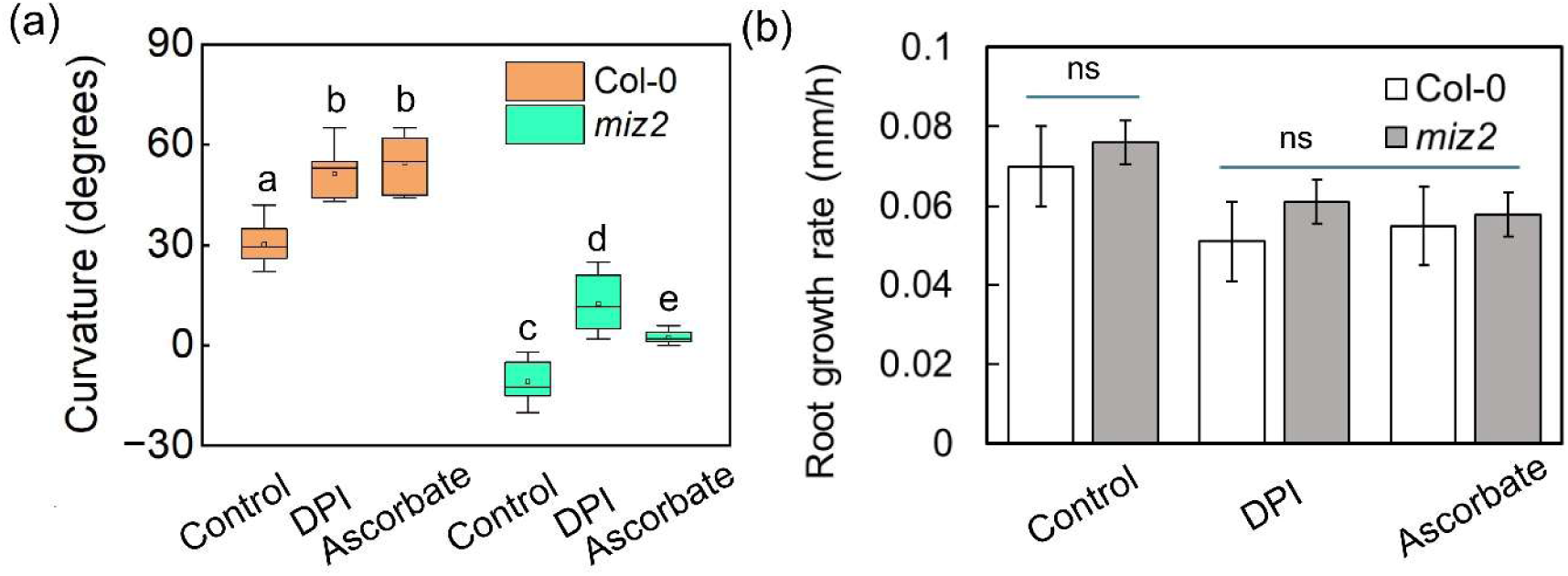
DPI and ascorbate affect *miz2* root halotropism. (a) Quantification of root-tip curvature in seven-day-old wild-type (Col-0) and *miz2* seedlings under control, 0.5 mM ascorbate, or 1 µM DPI treatments. Box plots represent the interquartile range; horizontal lines indicate the mean; whiskers denote SD (three independent experiments of 15 seedlings each; n = 45). (b) Root elongation rate calculated from primary root length before and after halostimulation in seedlings treated with control, ascorbate, or DPI. Values represent mean ± SD (three independent experiments of 15 seedlings each; *n* = 45); ns, no significant difference relative to Col-0 (Student’s *t*-test).

## Materials and Methods

### Halostimulation System under Continuous Clinorotation

To expose roots to a lateral NaCl gradient (halostimulation) while minimizing gravitropic influence, we adapted the split-agar method originally developed for hydrostimulation by replacing sorbitol with NaCl (Takahashi et al.,2002; Krieger *et al*. 2016; Yu *et al*. 2022). Five-or seven-day-old seedlings were positioned on control agar-solidified medium occupying half of a 90 mm Petri dish, with root tips placed 5 mm from the medium edge. The other half of the plate was filled with medium supplemented with NaCl (200 mM), allowing gradual diffusion toward the roots (Figure **S1**). Plates were sealed with parafilm and mounted on a benchtop rotator (ELMI Intelli-Mixer™ RM-2L) rotating at 9 RPM to achieve continuous clinorotation during the assay. For chemical treatments, seedlings were placed on medium containing ascorbic acid (0.5 mM in DDW) or diphenyleneiodonium (DPI; 1 μM in DMSO) 1 h prior to halostimulation. Halostimulation was then initiated by adding the NaCl-containing half of the split-agar system to the medium. Seedlings were imaged using a Nikon D7100 camera with an AF-S DX Micro NIKKOR 85 mm f/3.5G ED VR lens (Nikon, Tokyo, Japan), and root-tip curvature was quantified using ImageJ software (https://imagej.nih.gov/ij/).

### Confocal Microscopy and ROS Detection

ROS accumulation in roots was visualized using the H₂O₂-sensitive probe dihydrorhodamine 123 (DHR), as previously described (Gomes et al., 2005; Krieger et al., 2016). Briefly, immediately following halotropic stimulation, seedlings were immersed in 86.5 μM DHR (0.003% w/v; Sigma-Aldrich) prepared in phosphate-buffered saline (PBS, pH 7.4) for 2 or 5 min. After staining, seedlings were briefly rinsed in PBS to remove excess dye and mounted in PBS on glass slides. Confocal imaging was performed using a Leica SP8 laser-scanning confocal microscope (https://www.leica-microsystems.com/products/confocal-microscopes/p/leica-tcs-sp8) equipped with a 10× air objective. Imaging settings were kept constant across all samples to allow quantitative comparison. Excitation was at 488 nm (2% laser power), and emission was collected between 519 and 560 nm. Detector master gain was maintained between 670 and 720 (instrument-specific arbitrary units), with digital gain set to 1. Images were acquired using identical pinhole size, scan speed, and resolution parameters for all treatments. Fluorescence intensity was quantified as mean gray value using ImageJ/Fiji. For length measurements, ROS domains were defined manually using segmented region-of-interest tools, applying the same thresholding criteria across samples. Confocal images were pseudo-colored (RGB LUT) in ImageJ for visualization purposes only, without altering raw pixel values.

